# Susceptibility to multitasking in chronic stroke is associated to damage of the multiple demand system and leads to lateralized visuospatial deficits

**DOI:** 10.1101/2023.10.11.561866

**Authors:** Elvio Blini, Daniela D’Imperio, Zaira Romeo, Michele De Filippo De Grazia, Laura Passarini, Cristina Pilosio, Francesca Meneghello, Mario Bonato, Marco Zorzi

**Affiliations:** Department of General Psychology, University of Padova, Padua, Italy; Department of Neurosciences, Psychology, Drug Research and Child Health, University of Florence, Florence, Italy; IRCCS San Camillo Hospital, Venice, Italy; Padova Neuroscience Center (PNC), University of Padova, Padua, Italy

**Keywords:** Stroke, Multitasking, Multiple Demand system, Multivariate lesion-behavior mapping

## Abstract

Functional impairment after stroke is related to the amount of brain damage but there is no strict correspondence between lesion and magnitude of the deficit or its recovery. Theoretical constructs such as cognitive or brain reserve have been invoked as unspecific protective factors to explain this mismatch. Here, we consider the opposite point of view, that is the instances in which this protection is lost or overturned. Several studies have shown – in domains encompassing sensory, motor, and cognitive deficits – that paradigms in which the inherent processing limits of the brain are stressed (e.g., by introducing multitasking and attentional load), are indeed capable to unveil the presence of deficits that are otherwise missed. We administered a computerized multitasking paradigm to a sample of 46 consecutive patients with first-ever unilateral subacute to chronic stroke and no sign of lateralized spatial-attentional disorders according to established diagnostic tests. We then used cluster analysis to classify patients, in a purely data-driven manner, according to their multivariate pattern of performance across different conditions (e.g., single- vs dual-tasking, ipsi- vs. contra-lesional stimuli). This enabled us to identify, within a group of putatively spared patients, a cluster of individuals presenting with stark contralesional biases of spatial awareness exclusively in conditions of concurrent attentional load, i.e. a phenotype characterised by high susceptibility to multitasking. This construct was also captured by a latent factor obtained from principal component analysis, providing a continuous susceptibility index across the whole sample. In spite of similar lesion volume, patients in the high susceptibility cluster presented worse performance in activities of daily living and neuropsychological tests evaluating domain-general abilities spanning attention, executive functions, and reasoning. Multivariate predictive modeling based on lesions location and structural disconnections revealed distinctive correlates of high sensitivity to multitasking in the Multiple-Demand (MD) System, a network of frontal and fronto-parietal areas subserving domain general processes. Damage in these areas may critically interact with domain-specific ones (e.g., devoted to spatial attention), resulting in subtle, but significant difficulties for patients in everyday life situations. In conclusion, the construct of susceptibility to multitasking has the promising potential to provide us a better understanding of what marks the passage, after brain damage, to clinically visible deficits.

**Graphical Abstract:** 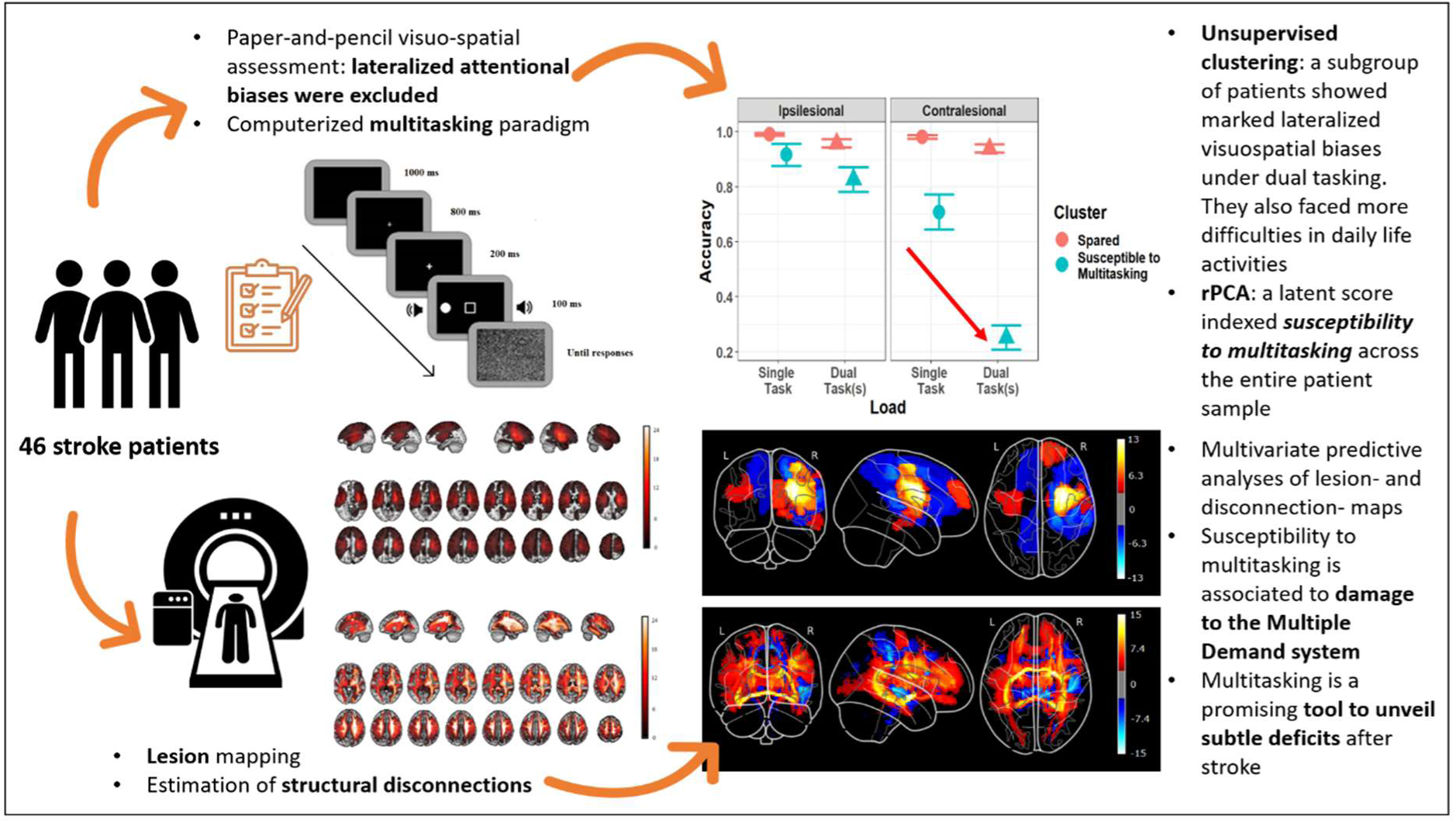

## Introduction

Stroke is one of the leading causes of death and morbidity worldwide^1^, with an estimated 50% of stroke survivors presenting significant sequelae. Neuropsychological research has typically focused on the characterization of (stark) behavioural impairments and the associated lesions to cortical and subcortical structures^2^, which is also very important for the study of recovery and rehabilitation^3,4^. The statistics imply, however, that a significant proportion of stroke survivors may, in fact, have minor consequences. Another positive news is that – despite the aging population, and a prevalence of stroke that is forecasted to sensibly increase in the western world^5,6^ – the advances in primary prevention strategies are gradually shifting the burden of stroke from mortality to disability. The widespread adoption of early treatments (e.g., thrombolysis) further contributes to containing this burden^6^. As a result, the line separating relatively spared patients and patients with some degree of impairment is shifting, but also becoming more blurred and uncertain. Therefore, it will be increasingly important, in the future, to study what marks the passage to stroke sequelae with enhanced resolution, as patients near this frontier will increase and may jeopardise the available clinical resources. The aim of this study is to better characterise where we should be drawing the thin, blurred line which, if crossed, determines the onset of cognitive deficits, with a special focus on disorders of spatial awareness.

The main path to a finer resolution in our assessment of patients’ deficits is developing tools that are sufficiently sensitive to a given construct (that is, a deficit). The most obvious way to achieve that is through technological advances. For example, when compared to clinical inspection, or tools that are designed for a coarse, quick clinical evaluation (for example, paper-and-pencil neuropsychological tests), computerized assessment offers a more sensitive and nuanced quantification of impairment and evaluation of rehabilitation outcomes^7–10^. A second, less obvious way, is informed by theory and is based on our state-of-the-art knowledge of brain functions and their inherent limitations.

Humans have, indeed, a limited capacity to process information^11^. Our performance typically declines when concurrent demands increase and/or serial information processing is required. For example, when a sequence of stimuli is rapidly presented in foveal^12^ or peripheral^13^ vision, the perception of a relevant target might be hampered if presented soon after the detection of a previous one, a phenomenon known as attentional blink. Concurrent task demands modulate the degree of interference by peripheral distracters^14^, suggesting that visuospatial abilities might be particularly hampered by multitasking. These well-known phenomena are comprehensively framed by the classic psychological literature that, since Broadbent’s seminal filter theory of attention, pinpoints the existence of bottlenecks in the flow of information in the brain^15^. The most evident obstacle is the case of multiple demands tapping onto the same sensory modality (e.g., visual tasks hampering visual perception). Multiple neuroimaging studies highlight, in the case of visual stimuli, occipital deactivation with increasing task demands^16,17^, which in turn suggests the existence of early peripheral processing bottlenecks that result in significant performance limitations. The behavioural impact of multitasking, however, extends beyond unisensory effects and can be domain-general or multisensory when higher-order processes are required. Working memory load, for example, similarly hampers visual search (to the point of inducing attentional blindness) and deactivates the temporo-parietal areas devoted to this main task^18–20^. These limitations are structural and computational in nature and, as such, can be consistently detected in the healthy brain. However, their impact is particularly stark and profound in the presence of a disturbance to said architecture, as in the case of a brain lesion affecting the optimal integration between brain networks.

Unsurprisingly, multitasking has been reported to exacerbate symptoms across behavioural domains, from more basic motor (e.g., balance and walking^21^, finger tapping^22^) or sensory functions (e.g., vestibular^23^), to higher level functions such as visual awareness^9,10,24^. In principle, multitasking can be added to any test, and it would presumably unveil even subtle deficits, if present, by creating a condition in which the available cognitive and attentional resources are diverted away and thus made unable to compensate for these difficulties^25^. Conversely, an increased susceptibility to multitasking caused by brain damage can be regarded as a major determinant of the emergence of frank deficits in a given behavioural domain whenever the context is more taxing due to concurrent task demands. Indeed, besides being transversal in nature (i.e., affecting diverse behavioural domains alike), multitasking conditions are also arguably ubiquitous in daily life situations, which better speaks to the generalizability of these measures, and may provide a hallmark signature of the outcome of a stroke.

This notion bears a striking resemblance with the concept of “strategy application disorder”, i.e. patients showing deficits restricted to conditions of multitasking following lesions of the right dorsolateral prefrontal cortex^26,27^. Action planning and task-switching have been classically considered a hallmark of executive functions, and their malfunctioning may very well subtend the emergence of multitasking deficits. However, a recent meta-analytic review suggests that multitasking and task-switching also present distinctive brain activation profiles^28^. While both dual-tasks and task-switching activate the bilateral intraparietal sulcus (IPS), left dorsal premotor cortex (dPMC), and right anterior insula, dual-tasking more specifically engages the bilateral frontal operculum, bilateral dPMC, bilateral anterior IPS, left inferior frontal sulcus, and left inferior frontal gyrus^28^. That said, the inconsistent use of labels and the different behavioural paradigms led to numerous discrepancies in the neurophysiological literature, pointing to the need of a unified framework to study dual-tasking^29^. The concept of multiple-demands system^30^ – describing a pattern of frontal and parietal activity associated with diverse cognitive tasks, as well as fluid intelligence – by stressing the commonality between different tasks and domains, is one potential candidate for this role. Susceptibility to multitasking might also be related to the notion of cognitive / brain reserve, which is typically invoked to explain the mismatch between the objective quantification of brain damage and the resulting clinical outcome^31–33^. In this regard, cognitive reserve is a major protective factor, shielding from cognitive deficits after brain disease, while susceptibility to multitasking might be regarded as the potential for this protective action to be overturned by unspecific demands. Therefore, quantifying susceptibility to multitasking, under these assumptions, enables the identification of the grey area in which brain damage can or cannot result in overt behavioural symptoms.

In this study, we capitalise on a dual-task paradigm originally proposed by Bonato and colleagues^24^ which, building on a purely top-down manipulation of task demands (that is, preserving invariance of physical stimulation), successfully highlighted subtle lateralized biases of spatial attention in patients with unilateral brain damage^8–10,24^. The task requires patients to report the side of appearance of one or two briefly presented dots that are displayed on the left, right, or on both sides of the screen, hence mimicking the classic diagnostic test for visual detection^34^. However, in all conditions, dots appear simultaneously to the presentation of a sound and a visual shape (see **Fig. 1**, for a graphical depiction). After providing a response to the primary spatial monitoring task (reporting dot side), patients are asked to either report the shape (dual-visual task, VDT), the sound (dual-auditory, ADT), or none of these additional features (single task, ST). Note that the single task does not require active encoding of information (beyond dot position) in order to be successfully performed, and it is therefore considered an active baseline condition (i.e., without attentional load). Previous small-scale studies on stroke patients^8,9^ have shown that multitasking conditions are strikingly more sensitive to lateralized deficits than single task conditions. In other words, the emergence of lateralized biases under dual-task conditions only is ideally suited to probe susceptibility to multitasking.

**Figure 1.**
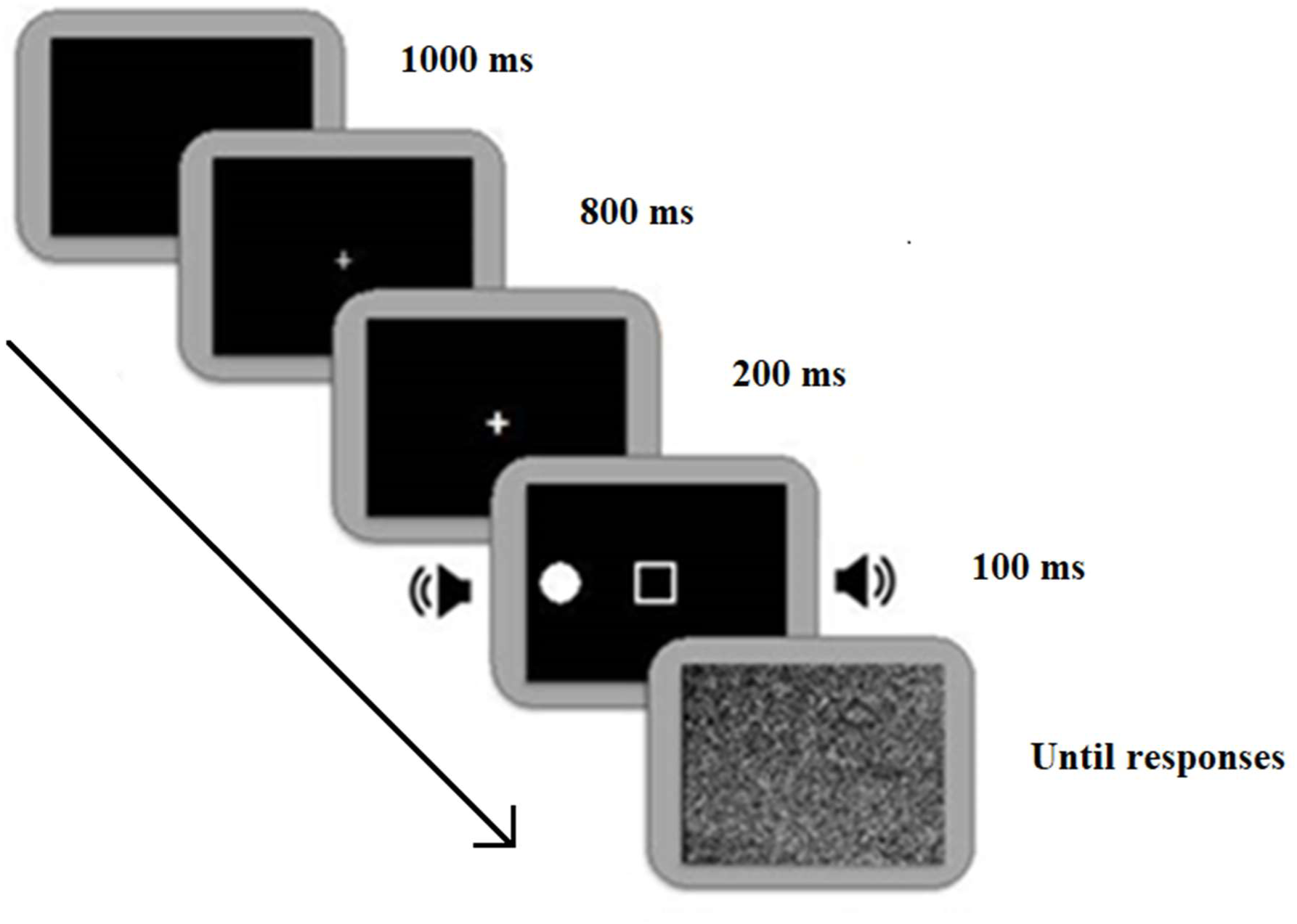
Schematic representation of the multitasking paradigm. The task consisted in a primary spatial monitoring task, in which the location of small dot(s) flashing briefly on the screen had to be reported (i.e., left, right, both sides). In the Single Task (ST), this was the only requirement. In the Auditory Dual Task (ADT), patients also had to report the image corresponding to the sound that they previously heard through headphones, whereas in the Visual Dual Task (VDT) they had to report the shape previously presented at fixation. The condition was made explicit in the instructions, which were given before each block. Note that conditions only differed for their task demands, but not for sensory properties. Thus, the difference in performance between single and dual-tasks is particularly suited to evaluate susceptibility to multitasking.

We administered the multitask paradigm to a sample of 46 consecutive patients with unilateral subacute or chronic stroke and no sign of lateralized attentional disorders according to established paper-and-pencil diagnostic tests (i.e., the Behavioral Inattention Test, BIT^35^), with the aim of identifying a phenotype characterised by high susceptibility to multitasking as well as its neuroanatomical correlates. First, K-means cluster analysis was used to classify patients, in a purely data-driven manner, according to their multivariate pattern of performance across conditions. This enabled us to identify, within a group of putatively spared patients, a cluster of individuals presenting with stark contralesional biases of spatial awareness only in the conditions of concurrent task demands. Moreover, Principal Component Analysis on the multivariate behavioural scores yielded a latent factor providing a finer-grained, continuous measure of this deficit. We then assessed the neuropsychological profile of these patients, as well as ecological measures of difficulties in everyday life situations. Finally, we investigated for the first time the anatomical correlates of susceptibility to multitasking using a multivariate predictive approach. Since behavioural deficits in stroke reflect both structural damage at the site of injury and more widespread network dysfunction caused by lesions to white-matter tracts, we examined (indirect) structural disconnection^36^ in addition to assessing the role of lesion size and location. In this regard, a crucial question was whether the multitasking-induced spatial awareness deficit is primarily related to damage affecting a domain-specific visuospatial mechanism, such as the right-hemisphere perisylvian network associated to hemispatial neglect^37^, or a domain-general mechanism such as the Multiple-Demand (MD) system^30^ involving the frontal regions that subtend dual-tasking^28^.

## Materials and methods

### Participants

Forty-six stroke patients (age: mean= 61.5y, SD=12.62, M= 33, F=13) in the subacute or chronic phase were included in this prospective study. The sample was composed of all the eligible patients among all those admitted to IRCCS San Camillo Hospital (Venice, Italy) for stroke rehabilitation in a time window spanning about 6 years (from March 2016 to September 2021). The enrolment of patients had the following inclusion criteria: adult age, first-ever unilateral stroke, right-handedness as assessed by a standard questionnaire^38^, and performance above the cut-off point in the conventional part of the Behavioural Inattention Test (BIT), that is the standard paper-and-pencil battery for the assessment of spatial neglect^39^. Exclusion criteria were the presence of a history of other neurological or psychiatric disorders, primary hearing, sight, or clinically evaluated visual field deficits, as well as the inability to understand and provide informed consent. The final consecutive sample included 29 patients with right hemisphere damage (RHD) and 17 patients with left hemisphere damage (LHD). All patients underwent comprehensive neuropsychological assessment and were administered the computerized task assessing susceptibility to multitasking. Structural brain scans (MRI, or in a few cases CT) were also collected for lesion analysis.

All participants gave their informed written consent, in accordance with the Declaration of Helsinki. The study protocol was approved by the regional Ethics Committee (Comitato Etico per la Sperimentazione Clinica della Provincia di Venezia e IRCCS San Camillo; protocol n. 2015.09 and n. 2018.04).

### Neuropsychological assessment

The neuropsychological assessment explored general cognition (Mini-Mental Scale Examination - MMSE^40^), reasoning (Raven’s progressive matrices^41^), selective attention (Attentional matrices test^42^), executive functions (Modified Card Sorting Test - MCST^43^), and memory (different tests according to the side of injury: for patients with RHD Rey’s auditory verbal learning test^44^; for patients with LHD Rey figure^45^). For visuo-spatial attention, we administered both the BIT^39^, as per our inclusion criteria, and the Kessler Foundation Neglect Assessment Process^46^, which provides a functional examination of daily living autonomy as impacted by spatial awareness deficits. The KF-NAP is built upon the Catherine Bergego Scale (CBS^47^); accordingly, few missing data for the KF-NAP were replaced by CBS scores in order to increase statistical power for this test (values were adapted to the appropriate range, i.e. 0- 10, with increasing values indexing more impaired performance). A language comprehension test was additionally administered to LHD patients to quantify the degree of linguistic impairments (Token test in the Aachener Aphasie Test – AAT^48^).

### Computerized task

We administered patients with a computerised multitasking paradigm previously optimised for patients with left and right hemisphere damage^9^. Patients sat in a quiet room, at about 60cm from the computer screen (38x30.5cm). The task ran through E-Prime 2.0 (Psychology Software Tools, Pennsylvania, USA, http://www.pstnet.com). Each trial started with a black screen (1000ms), followed by a centrally-presented white fixation cross (about 1cm wide, 1000 ms). The fixation cross flickered for 200ms before target presentation, as to direct attention to the screen centre. The target was a white dot (approximately 8mm in diameter), which appeared for about 100ms in three different locations: unilaterally, on the left or right side of the screen, or bilaterally at both sides of the screen (in all cases 170mm away from fixation). Additional “catch” trials were administered and did not involve any lateralized target. All trials were equiprobable (i.e., 25% of frequency). Simultaneously to the primary target(s), and for a duration of 100ms, a visual shape (i.e., square, circle or triangle) replaced the central fixation cross, and a sound (i.e., one of three ecological sounds, i.e. train whistle, hammer or doorbell) was binaurally delivered through headphones. Finally, a noise mask was presented until response, only after which the following trial could start (**Figure 1**).

The paradigm consists of three conditions: single task (ST), visual-dual task (VDT), and auditory dual-task (ADT). Task demands differed across conditions whereas sensory stimulation remained identical throughout the experiment. Patients were always asked to detect the position of the lateralized dots, i.e. spatial monitoring. In dual-task conditions, however, patients had to additionally report the central shape (VDT) or the sound (ADT). In case of difficulties in naming, patients could respond by pointing to a flashcard with a drawing of the possible options placed in front of them. The experimenter coded the responses on a keyboard and monitored the patient’s eye-gaze in order to discard trials in which fixation was not maintained.

The experimental paradigm consisted of a first ST practice block with 21 trials, followed by 6 experimental blocks, 36 trials each, administered in a fixed order (i.e., ST, VDT, ADT, VDT, ADT, ST), for a total duration of about 40 minutes.

## Behavioral data analysis

### Data preprocessing

Practice trials, used for familiarisation, were discarded. Catch trials, which were included to rule out guessing strategies or the presence of productive symptoms (e.g., perseverations), were also excluded based on the low proportion of false alarms (less than 6% on average) across both RHD and LHD patients. All trials in which the dot-target was presented (i.e., left, right, or both sides) were coded as ipsilesional, contralesional, and bilateral, in order to make LHD and RHD patients directly comparable. Accuracy of target detection for each patient was therefore computed as a function of Location (i.e., Ipsilesional, Contralesional, and Bilateral) and Load (i.e., ST, ADT, VDT).

### Patients’ clustering

In order to identify potentially distinct phenotypes based on performance in the computerized task, we used a purely data-driven (unsupervised) k-means clustering algorithm on the multivariate pattern of accuracy across the 9 variables given by the 3x3 combination of Location (Ipsilesional, Contralesional, and Bilateral) and Load (ST, ADT, VDT). The procedure identifies the best number of groups *k* which minimizes the within-groups variance (Euclidean distance) and maximizes the between-group variance. The resulting *k* groups, thus, are as distinct as possible but also internally consistent, such that the performance of patients belonging to the same cluster is similar whereas patients belonging to different clusters present substantial qualitative and/or quantitative differences. The best number of clusters *k* was chosen based on inspection of the scree plot depicting within-groups variance by *k*. However, the internal consistency of this parameter was also probed by repeating the clustering procedure with two subsets of the data. The two subsets represented “short” versions of the computerized task, in which only the first block of multitasking trials (i.e., either ADT or VDT) was assessed in conjunction with the first block of baseline trials (ST). Besides checking the internal reliability of the task and algorithm, these analyses also assessed the feasibility of a more time-efficient, brief version of the task which includes only one third of the trials. Finally, it is worth noting that the data-driven clustering procedure dispenses from the problem of establishing a somewhat arbitrary performance cutoff to assign patients to subgroups. It is worth noting that healthy elderly and even patients with Mild Cognitive Impairment tend to perform very close to the ceiling in our computerized task^9^, suggesting that it is not sensitive to unspecific, global cognitive impairments. Thus, k-means clustering allows us to detect patients with deviant performance within the overall sample of stroke patients.

Once *k* was chosen, we assessed whether RHD and LHD patients were equally represented in the clusters through a chi-squared test. Performance in the computerized task was assessed using analysis of variance (ANOVA) with Location and Load as within-subjects factors and cluster as between-subjects factor. Then, we assessed whether the newly defined clusters differed in terms of clinical variables (lesion volume, time from stroke onset, aetiology), demographic variables (age, education), and performance in conventional neuropsychological tests; Welch’s correction for unequal groups’ variances was used for t-tests when appropriate (this correction leads to degrees of freedom with decimal points).

### Data reduction: Principal Component Analysis

The patients’ multivariate patterns defined by the 9 accuracy variables (3x3 combination of Location and Load conditions) were also submitted to rotated Principal Component Analysis (rPCA) to summarize the behavioral performance with few latent factors. We used an oblique rotation (promax) because, for behavioural data, we favoured interpretability of the components as psychological constructs, and we did not want to force them to be orthogonal, as clinical factors may often correlate. We interpreted rPCs against overall performance and spatial biases in the computerized task. Moreover, we exploited the rPC(s) continuous values to explore the correlation with neuropsychological and neuroimaging data

## Neuroimaging data analysis

### Lesion analysis

Individual brain lesions were reconstructed from the patients’ MRI T1-weighted images (N=37) or CT scans (N=8). One patient did not give consent to be scanned. Each patient’s lesion was reconstructed from the original data and spatially registered on a common coordinate template provided by the Montreal Neurological Institute (MNI152 space, 91×109×91 with isovoxel resolution of 2mm) for further processing. For MRI scans, an automated segmentation of brain lesions was performed first using the Lesion Identification with Neighborhood Data Analysis software (LINDA^49^) and then, after visual inspection, manually corrected with the ITK-snap software^50^ by two experts (DD and ZR). Normalization into MNI152 space was performed using the pipeline of the BCBToolkit software^51^ (http://toolkit.bcblab.com), which is based on an enantiomorphic approach^52^ and uses affine and diffeomorphic deformations for image registration. Data from 5 patients had to be excluded from the analyses due to normalization failure. For CT scans, the lesion maps were manually segmented (DD and ZR) and then normalized using the RegLSM software, running on Matlab^53^. This method consists of two steps: firstly, native segmented CT is registered to an intermediated template, specific for elderly brains^54^, and then to the MNI152 template (the same space of MRI scans). Note that normalization to MNI152 is mandatory for computing structural disconnections (see below, also see Salvalaggio et al. ^36^ for discussion).

### Structural Disconnection

Structural disconnections were computed through the BCBToolkit software^51^ from each MNI-registered lesion map using 176 healthy controls from the “Human Connectome Project” 7T diffusion-weighted imaging dataset to track fibers passing through each lesioned voxel. The resulting structural disconnection (SDC) maps show only disconnected tracts and indicate the probable location of disconnections^55^, with each voxel reflecting the probability of disconnection (from 0.5 to 1 if above the conventional threshold of 0.5 and 0 otherwise^55^).

### Multivariate lesion and structural disconnection symptom mapping

Multivariate brain-behaviour mapping from lesions and disconnection maps was performed using the established approach of dimensionality reduction of the maps (each consisting of 902 229 2-mm^3^ brain voxels) through PCA followed by multiple regression onto a target behavioural score^36,56^. We retained from PCA the components that explained 95% of the variance (that is, 22 components for lesion and 23 components for SDC maps) and used them as input for the regression model. Regression modelling exploited a Best Subset Regression (BSR) strategy using, as dependent variable, the first component obtained from the rPCA on the computerized task scores (rPC1); rPC1 was multiplied by -1 prior to modelling in order to ease data interpretation, so that positive values indicated the presence of a deficit. BSR is a model selection procedure that consists in iteratively testing all possible combinations of predictor variables (with simultaneous inclusion of up to 10 predictors at each iteration). We selected the best model using the Bayesian Information Criterion (BIC), which is a stringent selection criterion that strongly penalizes model complexity (thereby minimizing the number of predictors in the final model). We thus obtained the best model for both lesion and disconnectome maps and assessed its cross-validated predictive accuracy in terms of R-squared. To control for the possible influence of age, lesion volume, and time from stroke onset, these variables were among the possible predictors in the BSR analysis. All covariates were scaled beforehand as to have unit-variance (e.g., direct total lesion volume control, dTLVC^57^).

Finally, the models’ regression weights were back-projected onto the original space using the transpose matrix of the PC coefficients^36^. The back-projected values were Z-normalized and smoothed with a Gaussian kernel (sigma = 1 mm) and thresholded at Z=3 to display the voxels that are most predictive of the behavioral deficit. Predictive maps were visualized in a glass brain (using the niilearn Python library) and the positive Z-scores (which are associated to deficit) were also rendered as a mosaic of 2D slices using MRIcroGL^58^ (http://www.nitrc.org/). An atlas-based approach^59^ was used to label the clusters of predictive voxels. For the lesion map we used the Automated Anatomical Labeling (AAL) atlas (version 3) of gray matter structures^60^. Furthermore, predictive white matter tracts were matched to a tract atlas^61^ for identification.

### Data availability

Computerized task code and predictive maps for lesions and structural disconnections are available at https://github.com/CCNL-UniPD/Multitasking. The conditions of our ethics approval do not permit the public archiving of the data. Additional summary data and code from this study are available upon reasonable request from the corresponding author.

## Results

Demographic, neurological and behavioral data about the whole patients’ sample are reported in **Table 1**. Other information about the complete neuropsychological patients’ profiles are reported in the supplementary materials (**Table S1**).

### Identification of clusters from the computerized task

Cluster analysis on the multivariate pattern of spatial monitoring accuracy yielded two patient clusters as the best solution. The first cluster (C1, N= 31) included 13 LHD and 18 RHD patients, that is 67% of the sample; the second cluster (C2, N= 15) included 33% of the patients’ sample, with 4 LHD and 11 RHD. The distribution of RHD and LHD patients did not differ between clusters (*X^2^*_(1, N= 46)_= 0.46, p = 0.496). We repeated the clustering procedure twice using only one block of ST and one block of dual tasking (either ADT or VDT), and checked the new predictions against the original clusters. Both brief versions yielded excellent consistency despite having only one third of the trials compared to the full task. In the brief-visual version (ST+VDT), three patients were misclassified. The balanced accuracy was 91.7%, significantly superior to the No-Information Rate (NIR) of 67.4% (binomial test: p<0.001, 95% CI [82.1%, 98.6%]). In the brief-auditory version (ST+ADT), only two patients were misclassified. The balanced accuracy was therefore 93.3%, significantly superior to the NIR (binomial test: p<0.001, 95% CI [85.2%, 99.5%]).

Overall, patients were accurate in reporting the central shape in the VDT (88.59%±13.05) or the sound in the ADT (92.12%±12) (i.e., the secondary task). The performance of patients in the primary visuospatial task is depicted in Figure 2 as a function of cluster. Notably, patients in C1 consistently achieved near-ceiling performance across all conditions. Patients in C2, however, showed a distinctive pattern of awareness deficit for contralesional targets (i.e., a neglect-like performance) only during multitasking. The 3 (Location) x 3 (Load) x 2 (Cluster) ANOVA on accuracy showed a significant three-way interaction, (F_(4,176)_= 4.07, p= 0.0035, pes= 0.085), indicating that the two clusters indeed differ in their susceptibility to multitasking. A finer-grain assessment is described below, in terms of different latent factors defining the two clusters. Overall, the results of the clustering procedure are very clear in pointing to the following conclusions:

1. Two distinct behavioral phenotypes are found when assessing performance under multitasking. One cluster (C1) presents an overall spared, and near-perfect performance, whereas a second cluster (C2) shows markedly compromised performance during multitasking.
2. The difficulties encountered by C2 patients are not generalized nor unspecific. Indeed, their performance in Ipsilesional trials was largely spared, with more than 83% of correct responses across Load conditions, and thus was not driving the assignment to a distinct cluster. The sharpest performance drops occurred in Contralesional and Bilateral trials, which is a marker of spatial awareness deficit for the contralesional side of space.
3. Crucially, a marked spatial awareness deficit emerged in C2 only when concurrent task-demands were introduced, irrespective of whether the concurrent task engaged the same or a different sensory modality (i.e., VDT vs. ADT). In these conditions, performance of C2 was as low as 22% of correctly-reported targets, whereas in the baseline (i.e., ST) condition the average accuracy remained above 62%. Thus, a context in which attentional resources are limited is particularly prone to unveil any underlying attentional bias, whereas standard conditions may not present sufficient sensitivity and may allow patients to fully compensate for this bias.

**Figure 2.**
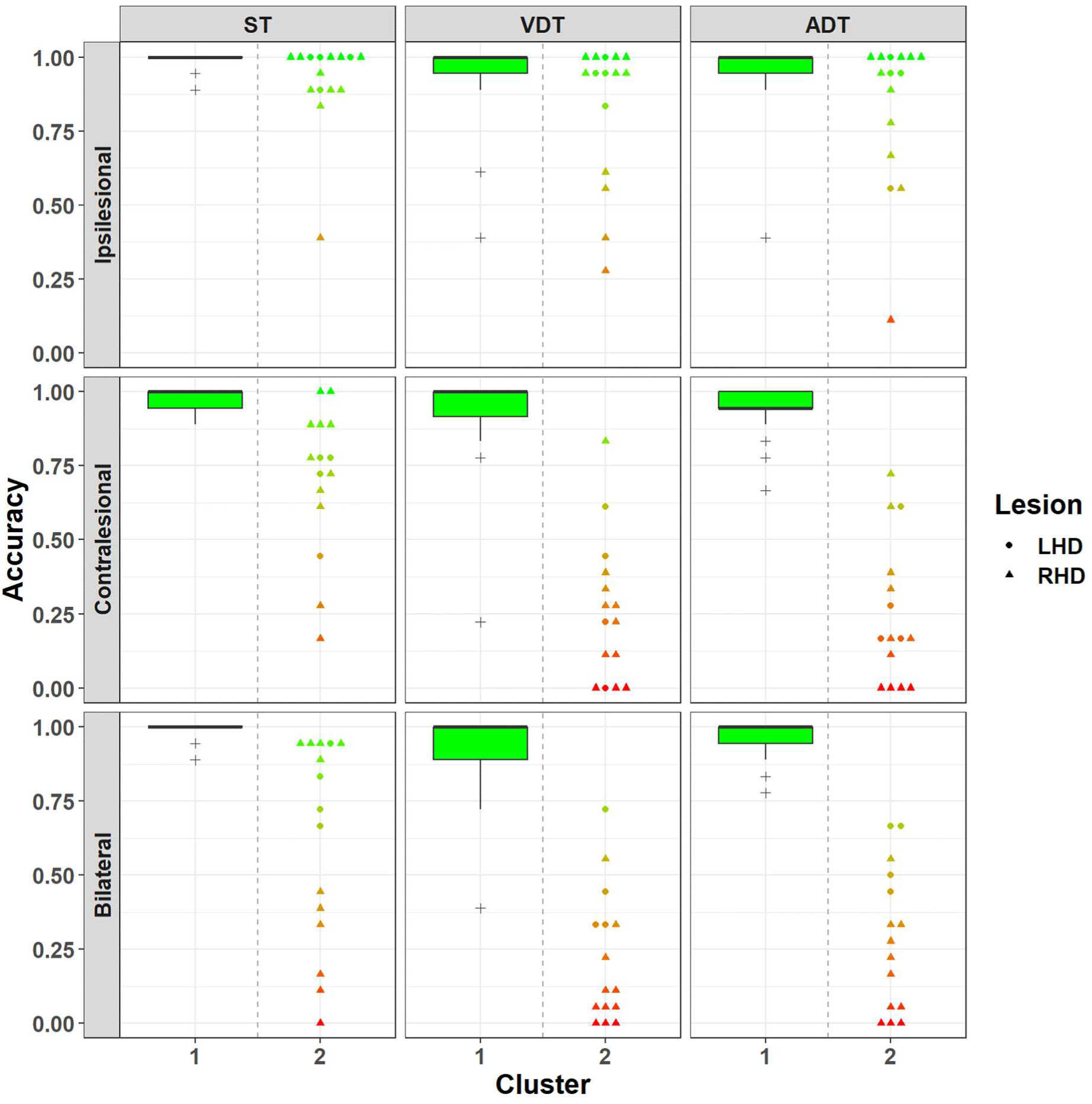
Patients’ clustering. Accuracy in the computerized task is depicted as a function of dot location (i.e., Ipsi-, Contra-, or Bi-lateral presentation) and load condition (i.e., Single Task - ST, Auditory Dual Task - ADT, and Visual Dual Task - VDT). An automated k-means clustering algorithm identified two clusters of patients within the entire sample (N= 46). Cluster 1 (C1, N= 31) achieved near-ceiling performance, whereas Cluster 2 (C2, N= 15) was characterized by clear lateralized biases which mainly emerged under multitasking conditions. Data for C1 is depicted as boxplots to avoid overcrowding, as this group was numerically larger and showed small variability. Individual patients in C2 are represented differently according to lesion side (circle: Left Hemisphere Damage - LHD; triangle: Right Hemisphere Damage - RHD).

### Control for lesion volume, time from stroke, age, and education

The two clusters did not differ in terms of overall lesion volume (t_(19.43)_= 1.09, p= 0.29; C1: 49.47±69.56cm^3^, C2: 79.79± 88.38cm^3^), neither for aetiology of lesions (*X^2^*_(1, N= 46)_= 0.72, p = 0.395). Time from stroke onset was also comparable in the two clusters (t_(29.47)_= 0.38, p= 0.70), as all patients were tested in the subacute and chronic stage (C1: 10.79±19.76 months,; C2: 9.06±18.52 months). However, patients in C1 were significantly younger (57.3±12.63 years) than patients in C2 (70.1± 7.31 years), t_(42.37)_= 4.31, p< 0.001, while there was no difference in terms of years of formal education (t_(26.82)_= 0.29, p= 0.77).

Age difference thus appears an important feature of C2, which is not entirely surprising considering that dual tasking performance (at least in terms of latency measures) declines in healthy ageing^62^. To control for the potential confound of age in the clustering solution, we iteratively removed the younger patients from C1 until reaching evidence that the two clusters were comparable in terms of age (using a Bayes Factor <1 for the contrast as criterion). This occurred after the exclusion of the 12 youngest patients in C1, yielding a subgroup of N= 19 patients with mean age of 65.68 years (SD= 7.79) that did not differ from the C2 patients’ mean age (t_(30.98)_= 1.69, p= 0.1; BF=0.95). We therefore repeated the 3*3*2 ANOVA (Load * Type * Cluster) with this new, age-matched, control group and found that the three-way interaction remained significant (F_(4,128)_= 2.62, p= 0.038, pes= 0.076) and with a similar effect size (see **Figure S1**, Supplementary Materials). Crucially, we also repeated the clustering procedure above without the 12 youngest (C1) patients and observed 100% concordance between the new cluster assignment and the one previously obtained on the full sample. Overall, these findings show that impaired contralesional performance under multitasking (and therefore assignment to the C2 cluster) is not driven by age, education, or clinical variables such as lesion volume and time from stroke.

### A continuous index of susceptibility to multitasking from rPCA

Three components in the rPCA solution accounted for 87.9% of the cumulative variance of the data (see components’ loadings in **Table S2** in supplementary materials). The first component (rPC1), in particular, explained 41.9% of the variance and loaded especially onto lateralized conditions (i.e., Contra- and Bi-lateral) but only under multitasking (i.e., both AVT and VDT), similarly to the main feature of C2. Indeed, we compared the values of this component between C1 (M= 0.65) and C2 (M= -1.34) to find that the two groups significantly differed along this dimension (t_(16.93)_= 15.43, p< 0.001). We therefore retained this component for all analyses warranting the use of continuous measures alongside the information about the clustering. The second (rPC2) and third (rPC3) components also accounted for an additional 23% of the variance each, and mostly loaded onto ipsilesional trials (rPC2) and trials belonging to the ST (rPC3). Thus, these last two components appeared to represent the “baseline” performance for the (main) effects of Type and Load. The three components were positively correlated, with values ranging from r= 0.32 to r= 0.58. Of these scores, only those of rPC3 significantly differed between C1 (mean= 0.40) and C2 (mean= -0.83) (t_(14.39)_= 3.38, p= 0.0043); note, however, that the correlation between rPC1 and rPC3 was r= 0.58, which indicates that the two constructs are somehow collinear and thus redundant. There was no difference between clusters in terms of rPC2 (t_(15.87)_= 1.59, p= 0.131).

### Relationship between susceptibility to multitasking and neuropsychological assessment

Group differences, as well as the correlation between each variable and rPC1, are reported in detail in **Table 2**. The two clusters did not differ in terms of MMSE scores (N= 41, t_(20.32)_= 0.965, p= 0.346). They differed, instead, in their average performance at the neuropsychological tests assessing attention: C2 presented marginally lower scores with respect to C1 in the BIT (C2: 138.1±4.27, C1: 141.9±2.75) as well as in the Attentional Matrices test (C2: 37.13±9.8, C1: 44.2±10.1) that specifically assesses selective attention. However, all patients had scores within normal range in both tests; thus, multitasking revealed an otherwise invisible spatial attentional deficit. Most importantly, the susceptibility to multitasking shown by C2 patients is not a mere subtlety because the two clusters significantly differed in terms of functional evaluation of daily living activities. Indeed, C2 patients presented significantly higher scores (indicative of more severe deficits) in the KF-NAP^46^, assessing spatial neglect-like biases and difficulties in ecological situations, compared to C1 patients (C1: 1.85 ±2.66, C2: 5.54±4.8; N= 40, *t*_(15.63)_= 2.58, *p*= 0.02).

All these neuropsychological tests were also significantly correlated with the first component obtained from the rPCA (though to a lesser extent for attentional matrices). Finally, although cluster differences were comparatively smaller in this case, we found that rPC1 correlated with the scores of two tests evaluating reasoning and executive functions, i.e. the Raven’s progressive matrices, and the Modified Card Sorting Test, the latter for both category scores and, marginally, for perseveration errors.

### Multivariate analyses of lesion- and disconnection-symptom mapping

The majority of lesions, for both LHD and RHD patients, affected the insular, central opercular cortex, and putamen. The most affected tracts were the superior longitudinal fasciculus III (SLFIII), fronto striatal fasciculus, pons and corpus callosum for both LHD and RHD patients. In addition, the corticospinal tract appeared frequently disconnected by the right hemisphere lesions (see **Figure S2** in the Supplementary Materials).

Multivariate brain-behaviour mapping was carried out both on lesion and SDC maps (N= 40 patients, 27 for C1 and 13 for C2). We first assessed the potential role of covariates such as lesion volume, time from stroke onset, and age. Neither lesion volume (r= -0.2, t_(38)_= 1.26, p= 0.22) nor time from stroke onset (r= -0.11, t_(38)_= -0.66, p= 0.52) were significantly correlated with rPC1, and did not differ between clusters (t_(19.43)_= 1.09, p= 0.29 for lesion volume; t_(29.47)_= 0.38, p= 0.70 for time from stroke). This is remarkable in that lesion size has been classically considered a proxy for stroke severity and has been often found to correlate with behavioral deficits, though in interaction with mediator variables^33,63^. Age, on the other hand, was significantly associated with rPC1 (r= -0.46, t_(38)_= -3.48, p= 0.001) and did differ between clusters, as outlined in the paragraph above. Thus, these three variables were included in the BSR approach as predictors alongside the features obtained from the neuroimaging data (22 for lesion maps, 23 for SDC).

### Lesion location

For lesion maps, the BSR approach selected a model including k = 5 predictors (PCs 3, 5, 7, 8 and 9), in addition to age and lesion volume as covariates (BIC= 104.4). This model was far superior to the null model (intercept-only: BIC=119.3). Note that smaller values index better fit and a BIC difference of 10 corresponds to a posterior odds of about 150:1^64^. When assessed using leave-one-out crossvalidation, the model’s predictive accuracy on left-out patients in terms of explained variance was R^2^=0.39, which is well aligned with state-of-the-art multivariate lesion-symptom mapping across behavioural domains^36^. When age and lesion volume were not in the pool of possible predictors, the modelling approach still selected k= 5 predictors (components 1, 5, 8, 9, and 14) and obtained a better fit than the null model (BIC= 109.6). Importantly, the performance in crossvalidation was very similar to that obtained with additional covariates (R^2^=0.36), showing that age alone was not driving the good model fit and performance.

Back-projection of the model coefficients to the atlas space for visualization (see **Fig. 3**) revealed that the right hemisphere voxels most predictive of susceptibility to multitasking corresponded to the frontal white matter (Z peak at [30,-4, 34]) and extended into the inferior frontal operculum, the insula, and precentral gyrus. This region included the most predictive voxels overall and had a symmetric counterpart (though with much smaller coefficients) in the left hemisphere ([-26, -6, 30]). Secondary clusters, associated with much smaller coefficients, were located in the thalamus (ventral lateral thalamus, [16,-12,0]), and the medial aspect of the right superior frontal gyrus ([10, 56, 28]).

**Figure 3:**
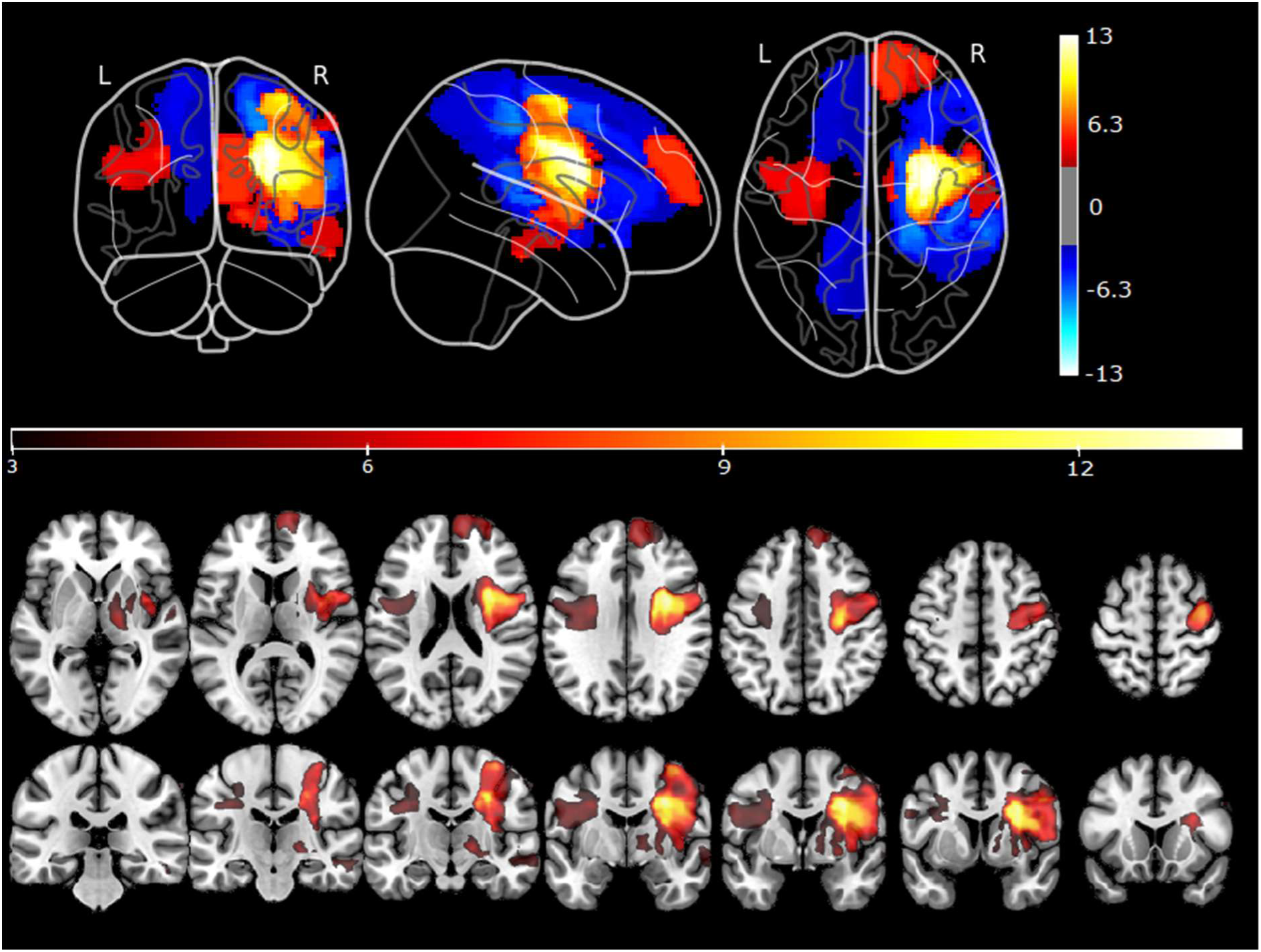
Predictive weights back-projected on the brain showing the association between susceptibility to multitasking (indexed by rPC1) and lesions. Top panel: Glass brain projection, where red-yellow represents voxels predicting deficits while blue-green represents voxels predicting no deficit. To optimize the visualization, Z-normalized values in the range (- 3, 3) are not displayed. Bottom panel: Back-projected values associated with the deficit (Z>3 for better visualization) are rendered on a mosaic of 2D slices.

Overall, predictive voxels were generally located more anteriorly with respect to the brain areas more typically associated with spatial neglect. The locations of predictive lesions were, on the other hand, much more in line with the involvement of a domain-general mechanism. In particular, the MD system has been previously associated with areas in the posterior part of the inferior frontal sulcus, in particular the frontal operculum and anterior insula^30,65^. Brain activations in the MD system have been associated with diverse cognitive demands and the assembly of many distinct subtasks^30^. Thus, one possibility is that impairments in this domain-general network may interact with or affect distant areas, among which those devoted to domain-specific (e.g., visuospatial) processes. This is in large agreement with recent evidence^66^ suggesting that disrupted anatomical connectivity of the MD system (as measured, for example, through indirect disconnections and as opposed to functional connectivity, measured with resting-state fMRI) predicted very well deficits in a range of diverse neuropsychological tests (e.g., the Trail-Making Test or the Stroop test), measuring different constructs but yet joined in their requiring a certain degree of cognitive control.

### Structural disconnections

For SDC maps, the BSR approach selected a model with k = 3 predictors (PCs 2, 3 and 20) and age as a covariate (BIC= 104.7), which was superior to the null model (intercept-only; BIC = 119.3). The crossvalidated predictive accuracy on left-out patients was R^2^= 0.39. When age was not in the pool, performance in cross-validation was slightly worse (R^2^= 0.24), although in face of a relatively good model fit with k= 3 predictors (PCs 1, 2, and 20; BIC= 112.4). The predictive map obtained by back-projecting the model coefficients is displayed in **Figure 4**. Susceptibility to multitasking was associated with widespread damage to the right and left anterior thalamic radiation (ATR) and right SLF II. In addition, the analysis also highlighted an important role of interhemispheric disconnection, in that predictive voxels extended to the opposite hemisphere through substantial involvement of the corpus callosum and frontal commissure. Note that the ATR has been previously linked to individual differences in multitasking ability in healthy individuals^67^. SLF II has been implicated in spatial awareness before^68,69^ as, if damaged, it predicts well the presence of spatial neglect. The latter disconnection seems to reconcile with the fact that lesion location does not suggest the involvement of domain-specific areas such as those subserving visuo-spatial attention, which might nonetheless become dysfunctional due to their connectivity with regions in frontal areas subserving domain general processes that are relevant for multitasking or cognitive control^66^.

**Figure 4.**
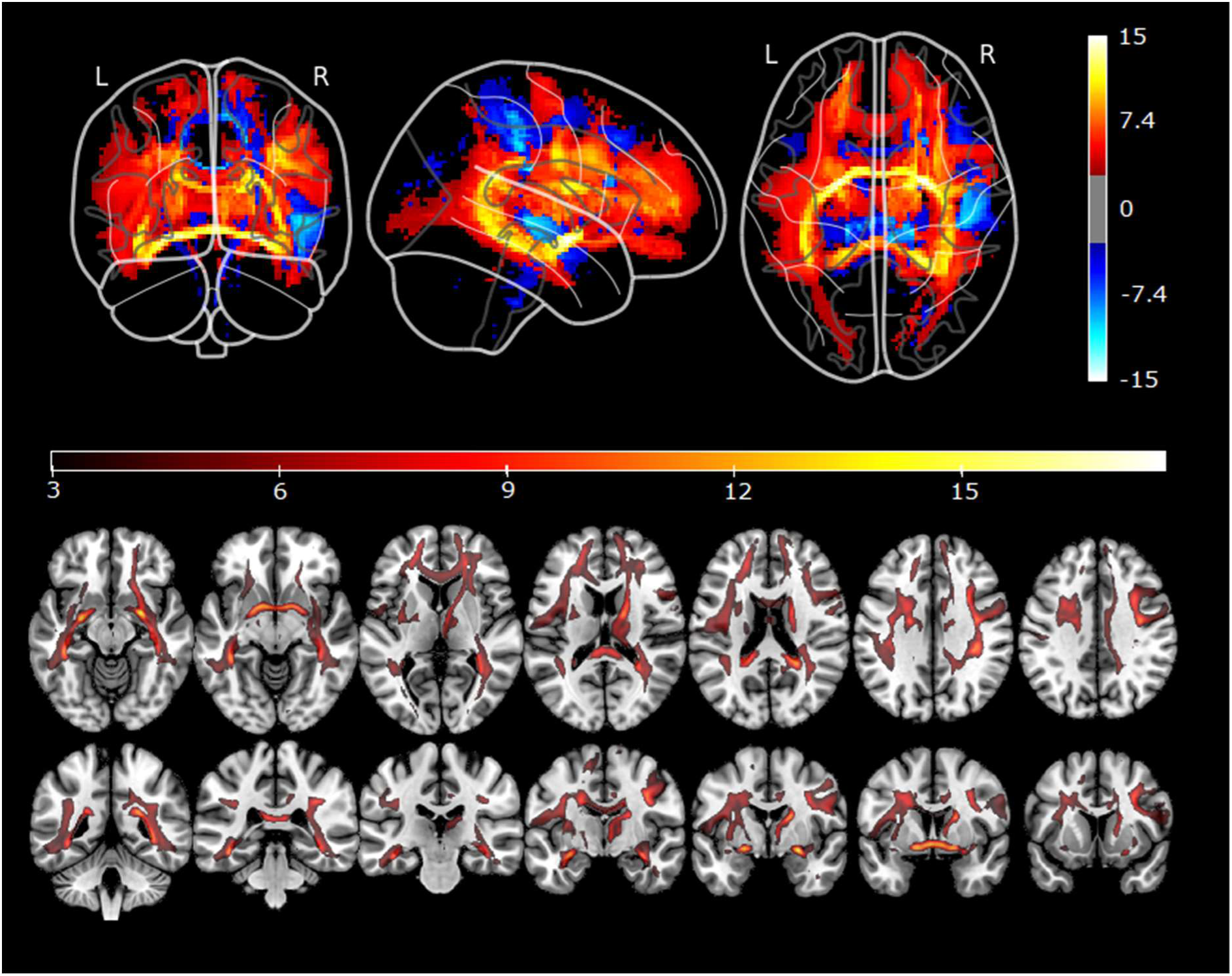
Predictive weights back-projected on the brain showing the association between susceptibility to multitasking (indexed by rPC1) and structural disconnections. Top panel: Glass brain projection, where red-yellow represents tracts predicting deficits while blue-green represents tracts predicting no deficit. To optimize the visualization, Z-normalized values in the range (−3, 3) are not displayed. Bottom panel: Back-projected values associated with the deficit (Z>3 for better visualization) are rendered on a mosaic of 2D slices.

## Discussion

In this study, we enrolled patients with first-ever subacute or chronic stroke who performed well within normal limits in a classic paper-and-pencil neuropsychological test used for the assessment of visuospatial neglect^39^. When administered with a computerized task tapping onto spatial monitoring performance, both in isolation and under conditions of concurrent task demands (i.e., multitasking, or increased attentional load), patients split into two very distinct behavioural phenotypes. Two-thirds of the patients (67%) performed the spatial monitoring task with near-ceiling accuracy, irrespective of target side (ipsilesional vs contralesional) and of the increasing attentional load in the multitasking condition. The remaining third of the patients (33%), however, deviated so markedly from the latter pattern that they were automatically assigned to a separate group by a multivariate clustering algorithm. This second cluster of patients (C2) was indeed impaired in the spatial monitoring task, but not in a generalized and unspecific way. In this group, performance in ipsilesional trials was always near-ceiling and even the detection of contralesional targets (for contralesional and bilateral trials) was relatively spared when spatial monitoring was the only task (i.e., the baseline, single task condition). Performance deteriorated sharply only when concurrent task demands were introduced, irrespective of whether the second task loaded onto the same (i.e., VDT) or a different sensory modality (i.e., ADT).

The finding of a behavioural phenotype characterized by high susceptibility to multitasking substantiates and qualifies previous small-scale studies suggesting that computerized tasks, especially when involving dual-tasking, can unveil subtle lateralized deficits of visuospatial attention^8–10^. A context in which attentional resources are limited is particularly prone to reveal any underlying bias, if present, whereas standard conditions may not have sufficient sensitivity and may allow patients to fully compensate for it^7^. Successful compensation for a deficit is a desirable feature, and may indicate that sufficient cognitive resources can be recruited when needed to overcome everyday life challenges and difficulties. But this condition may be fragile, as for a substantial share of patients these resources may be limited and rigidly capped. Hence, assessing susceptibility to multitasking allows one to objectively quantify the extent by which these patients would struggle in more taxing conditions, thereby enhancing our understanding of their needs. In other words, it enables us to better characterize what marks the passage to stark stroke’s sequelae, namely the buffer zone in which brain damage can or cannot result in significant deficits, depending on a wealth of contextual factors. Finally, it may inform treatment, in that patients within this gray area may benefit the most from cognitive training aimed at fostering general cognitive resources.

Our main task leverages on a classic visual detection paradigm for spatial neglect^34^, i.e. the inability to orient attention toward or respond to stimuli presented in the contralesional hemisphere^70,71^. However, it would be questionable to classify C2 patients as having spatial neglect, first and foremost because they all performed well above the clinical cutoff in established diagnostic tests (i.e., the BIT^39^). Traditionally, based on the results of clinical tests, all patients (both C1 and C2) would be considered as a control group (i.e., N-) for patients with overt lateralized spatial deficits (i.e., N+). It is also true, however, that C2 presents many core features of spatial neglect, though on a different scale. For example, spatial neglect is considered a rather heterogeneous syndrome in which core lateralized (i.e., spatial) biases coexist and interact with non-lateralized deficits^72^. This picture is very well captured by patients in C2: they show a marked lateralized bias (i.e., the difference in performance between Ipsilesional and Contralesional/Bilateral trials) when concurrently engaged by non-spatial demands. This suggests that the difference between C2 and patients with clinically-confirmed spatial neglect may be quantitative rather than qualitative in nature, which in turn has important consequences. First, equating the two clusters as both “spared” control groups may lead to a loss of potentially relevant information. In particular, neuropsychological and neuroimaging studies may overlook the clinical picture and neural correlates of patients in C2, thus failing to capture the full extent of spatial neglect manifestations. Second, an in-depth description of C2 may provide hints about the clinical characteristics of patients who are at the prodromal boundary of spatial awareness disorders, and possibly any other deficit, given that attentional load similarly affects several diverse domains^22,23^. This is not trivial because, as a matter of fact, patients in C2 may experience higher than normal functional difficulties in everyday life situations, as suggested by the higher scores (i.e., worse performance) of this group in the test evaluating activities of daily living (i.e., KF-NAP^46^). Thus, susceptibility to multitasking better captures the difficulties that may arise in everyday life, ecological situations.

One of the intriguing results of this study is that high susceptibility to multitasking was not correlated with the size of the brain lesion. Lesion volume is an established proxy for stroke severity, and it is generally assumed to correlate with virtually any deficit, although in interaction with mediator variables^33,63^. The lack of correlation with the main deficits of patients in C2 may question that they stem from a unitary construct. There was, on the other hand, an association between the latent factor indexing susceptibility to multitasking (rPC1) and patients’ age, which has been often taken, instead, as a proxy for brain reserve, chiefly because it is negatively associated with brain plasticity^33^. Older patients were more likely to present high susceptibility to multitasking, even in spite of similar lesion size. Thus, while patients’ difficulties may be graded and continuous in nature, this has not a continuous counterpart in terms of objective brain damage, at least in this study. While the quantity of brain damage does not seem to play a substantial role, multivariate lesion-symptom mapping linked susceptibility to multitasking to specific lesion locations and structural disconnections. We start by noting, first, that we did not observe a significantly different prevalence of high susceptibility to multitasking in patients with left (LHD) vs. right hemisphere damage (RHD). This may be surprising, in that disorders of spatial awareness are thought to be more common following lesion of the right hemisphere^73^, on the one hand, and because cognitive reserve may also be lateralized in the right hemisphere^31^, on the other hand. Moreover, as pointed out by De Renzi (1982)^74^, studies in which LHD and RHD are compared typically suffer a common selection bias: if only LHD patients with sufficiently spared language abilities are enrolled, in order for them to properly understand the experimental tasks, these patients will necessarily present, on average, milder stroke sequelae, and will generally perform better. However, we did not observe differences in this direction in the current study. Part of the reason is that our computerized task can be administered also to a population of patients with variable degree of language impairments^9^. Another possible explanation for this finding may be insufficient statistical power due to the different size of the groups (17 LHD vs. 29 RHD). Besides hemispheric asymmetries, however, multivariate lesion-symptom mapping did highlight the neural correlates of high susceptibility to multitasking in both hemispheres, suggesting that this construct may be differently lateralized than spatial attention.

The areas most predictive of susceptibility to multitasking were located in the frontal white matter, both for RHD and LHD patients, and extended to the frontal operculum and insula. Thus, the lesions associated with visuospatial bias under multitasking (i.e., the pattern shown by C2 patients) did not match those typically associated with spatial neglect^75^. This is not entirely surprising. First, while differences between C2 and clinically-confirmed spatial neglect may be only quantitative in nature, more extended lesions reaching, posteriorly, the inferior parietal lobe may be more likely to cause stark symptoms. There were no stark symptoms in our sample of patients, as per our selection criteria. Second, C2-like patients are likely to have been included in the control groups for studies on neglect, which may have led to overlook these frontal sites in most lesion-symptom mapping studies. The bilateral frontal operculum and anterior insula (together with bilateral dPMC, bilateral anterior IPS, left inferior frontal sulcus, and left inferior frontal gyrus) characterize the distinctive activation profile of dual tasking according to a recent meta-analysis^28^. These areas have been previously associated with several different tasks, spanning diverse cognitive functions, and may thus be regarded as an abstract, higher order, and multi-purpose system, known as the Multiple Demand network^30^, subtending overall cognitive functioning. For example, this network may provide the basis for cognitive control^66^, or the assembly and timing of specific subtasks^30,67^, all aspects that are independent, but closely related to the concept of fluid intelligence^67^. The MD system has been previously described in subcortical and cortical parietal and frontal areas, including the frontal operculum and the anterior insula^65^, thereby closely mirroring what was predictive of susceptibility to multitasking in the present study. It is therefore possible that structural damage to these regions would affect multitasking conditions more notably, which is precisely the main behavioural feature of C2. While this can nicely explain pure multitasking deficits, however, it still does not account for the full picture consisting of both a multitasking deficit and a concurrent spatial one. In this regard, the analysis of structural disconnections provides complementary and crucial information. White matter damage involving the ATR, anterior commissure, as well as fronto-parietal disconnections along SLF II were the most predictive of behavioral deficits. The ATR joins cortical (prefrontal) and subcortical areas, most notably the hippocampal formation, thalamus, and striatum^76^, subserving thus multiple tasks and functions^67,77,78^. Few studies have implied a role of the ATR in strategic processes, overall processing speed, and/or executive functions^78^; Kievit et al. (2014) found that the ATR integrity predicted age-related differences in multitasking ability. Overall, the ATR presents substantial connectivity with the prefrontal components of the MD system^67^, and thus its involvement is not entirely surprising given the most predictive lesions of susceptibility to multitasking, i.e. within the MD system itself. SLF II, on the other hand, is known to be implicated in spatial awareness^68,69^ as, if damaged, it predicts the presence of spatial neglect. SLF II originates from the posterolateral parietal lobe, including the angular gyrus, runs through the middle frontal gyrus, and ends within the dorsolateral prefrontal cortex^79^. Thus, the analysis of structural disconnections seems to reconcile the fact that the lesions of C2 were located in the frontal cortex, as opposed to areas more strongly tied to spatial processes such as those in the inferior parietal lobe: suboptimal communication between them caused by structural disconnection, which cannot be fully captured by lesion maps – may be ultimately responsible for the detrimental effects of multitasking on spatial attention. This would be in line with a recent study showing that lesions affecting the connectivity within the MD system, similar to what described here, lead to impairments in multiple aspects of attention that are traditionally considered relatively independent (e.g., visuo-spatial attention vs. alertness or executive control), corroborating the notion that these aspects closely interact and that a higher order factor may be the joining link^80^.

One important point to stress, however, is that damage to the MD system, albeit pervasive, should not be necessarily regarded as an alternative definition of “global impairment”. For example, the performance in ipsilesional trials of C2 patients was, even under multitasking, rather good, their deficits being limited to few critical conditions as defined by their underlying spatial awareness bias (contralesional or bilateral trials). Selective impairments, i.e. limited to peculiar domains or conditions, require a different explanation than a global deficit, and probably specific lesion patterns. One possibility is that the presence of an underlying “dormant” deficit may be necessary, as the disruption of the communication between candidate areas (one domain-general, one domain-specific). For this, future studies should probe the generalizability of susceptibility to multitasking to different measures (i.e., beyond spatial awareness).

In summary, computerized paradigms exploiting multitasking can detect subtle deficits even in seemingly spared, chronic stroke patients. These paradigms leverage on our state-of-the-art knowledge of the brain’s functions and limitations, and can be reliably administered even in very short, quick formats (e.g., 10 minutes). More importantly, they better capture the difficulties of patients in daily life activities, where multitasking is ubiquitous. Individual susceptibility to multitasking as a construct has the promising potential to provide us a better understanding of what marks the passage, after brain damage, to clinically visible deficits, hence advancing our knowledge of the impact of stroke and tailoring patient-specific options for treatment.

## Acknowledgements

We are grateful to all the patients who took part in our study. We would also like to thank the clinical team at IRCCS San Camillo Hospital for assistance and support.

## Author Contribution

**Elvio Blini**: Conceptualization, Methodology, Software, Formal Analysis, Data Curation, Writing – Original draft, Writing – Review and Editing, Visualization

**Daniela d’Imperio**: Conceptualization, Methodology, Investigation, Data Curation, Writing – Original draft, Writing – Review and Editing, Project Administration

**Zaira Romeo**: Conceptualization, Methodology, Investigation, Data Curation, Writing – Review and Editing, Project Administration

**Michele de Filippo de Grazia**: Software, Data Curation, Writing – Review and Editing

**Laura Passarini**: Resources, Writing – Review and Editing

**Cristina Pilosio**: Resources, Writing – Review and Editing

**Francesca Meneghello**: Resources, Writing – Review and Editing, Supervision, Project Administration

**Mario Bonato**: Conceptualization, Methodology, Writing – Review and Editing

**Marco Zorzi**: Conceptualization, Methodology, Formal Analysis, Resources, Data Curation, Writing – Review and Editing, Visualization, Supervision, Project Administration, Funding Acquisition

