## Supplementary information for "Susceptibility to multitasking in chronic stroke is associated to damage of the multiple demand system and leads to lateralized visuospatial deficits"

**Author affiliations:**

Full address: Department of General Psychology, University of Padova

Via Venezia, 12 - 35131 Padova - Italy

**Running title**: Susceptibility to multitasking in stroke

**Keywords:** Stroke; Multitasking; Multiple Demand system; Multivariate lesion-behavior mapping

**Supplementary Information**

**Behavioural data**

***Neuropsychological tests***

Scores from neuropsychological tests, in addition to those reported in Table 1 in the main text, are reported for the whole sample in Table S1.

**Table S1.** Scores of additional neuropsychological tests administered to patients.

| **Patient/Group** | **MMSE** | **Att. Matr.** | **KF-NAP** | **Raven** | **MCST**  **Cat.** | **MCST**  **Pers. err.** |  | **Memory** | **Token** |
| --- | --- | --- | --- | --- | --- | --- | --- | --- | --- |
| 1/RHD | 27 | 53 |  | 32 | 5 | 3 |  | 4 |  |
| 2/RHD | 30 | 49 | **2** | 30 | 6 | 1 |  | 3 |  |
| 3/RHD | **24** | 35 | **6** | 16 | **2** | 17 |  | 4 |  |
| 4/RHD | 30 | 38 | **9** | 28 | 5 | 5 |  | 4 |  |
| 5/RHD | 30 | 42 | **3** | 31 | 6 | 3 |  | 4 |  |
| 6/RHD | 29 | 25 | **1** | 24 | 3 | 11 |  | 4 |  |
| 7/RHD | 25 | 39 | **4** | 25 | **2** | 15 |  | 3 |  |
| 8/RHD | 29 | 53 |  | 30 | 6 | **0** |  | 2 |  |
| 9/RHD | 28 | 54 | 0 | 33 | 3 | 15 |  | **0** |  |
| 10/RHD | 30 | 47 | **3** | 29 | 6 | 3 |  | 3 |  |
| 11/RHD | 27 | **36** | **7** | 29 | 6 | 3 |  | 4 |  |
| 12/RHD | 28 | 52 | 0 | 26 | 5 | 8 |  | **0** |  |
| 13/RHD | 30 | 33 | **8** | 31 | 6 | 2 |  | 2 |  |
| 14/RHD | **26** | 40 | **15** |  | 5 | 2 |  |  |  |
| 15/RHD | 30 | 51 | 0 | 32 | 6 | **4** |  | 4 |  |
| 16/RHD | 29 | 53 | 0 | 33 | 6 | 4 |  | 3 |  |
| 17/RHD | 27 | 57 | 0 | 33 | 4 | 6 |  | 3 |  |
| 18/RHD | 26 | 56 | 0 | 34 | 4 | 6 |  | 4 |  |
| 19/RHD | 29 | 43 | **7** | 28 | **0** | 16 |  |  |  |
| 20/RHD | 29 | 40 | **6** | 29 | 6 | 3 |  | 4 |  |
| 21/RHD | 30 | 53 |  | 36 | 6 | 6 |  | 4 |  |
| 22/RHD | 27 | 36 |  | 29 | 3 | 13 |  | 4 |  |
| 23/RHD | 30 | 52 | **8** | 35 | 6 | 1 |  | 4 |  |
| 24/RHD | 30 | 50 | **4** | 33 | 5 | 8 |  | 4 |  |
| 25/RHD | 30 | 43 | **6** | 28 | 6 | 3 |  | 4 |  |
| 26/RHD | 30 | 48 | **6** | 27 |  |  |  | 4 |  |
| 27/RHD | 29 | 51 | 0 | 35 | 6 | **0** |  | 4 |  |
| 28/RHD | 27 | 52 | **13** | 23 | 6 | 5 |  | 4 |  |
| 29/RHD | 28 | 33 | **4** | 23 | **1** | 11 |  | 2 |  |
| 1/LHD | 29 | 48 | **10** | 36 | **2** | 2 |  | 4 | 2 |
| 2/LHD | **26** | 42 | 0 | 33 | **2** | **8** |  | 3 | 5 |
| 3/LHD | 26 | **34** | 0 | 34 | 4 | **13** |  | 3 | 22 |
| 4/LHD |  | 36 | 0 | 31 | **2** | **11** |  | 2 | 13 |
| 5/LHD | 30 | 54 | 0 | 27 | 4 | 11 |  |  |  |
| 6/LHD | 26 | 37 | 0 | 33 | 6 | 2 |  | 4 | 38 |
| 7/LHD | 27 | 50 | 0 | 24 | 5 | 4 |  | 3 |  |
| 8/LHD | 27 | **25** | **1** | 30 | **2** | **13** |  | 4 | 23 |
| 9/LHD | 28 | 40 |  | 22 | 5 | **9** |  | 4 | 17 |
| 10/LHD | 29 | 39 | 5 | 35 | 6 | 4 |  | 4 | 27 |
| 11/LHD | **26** | **33** |  | 34 | 6 | 0 |  | 1 | 1 |
| 12/LHD |  | **30** | **1** | 33 | **3** | **13** |  | 1 | 22 |
| 13/LHD | 27 | **19** | 0 | 28 | 4 | 7 |  | **0** | 8 |
| 14/LHD |  | **12** | 0 | 24 |  |  |  |  | 40 |
| 15/LHD |  | **30** | 0 | 34 | 3 | **7** |  | 4 |  |
| 16/LHD |  | **34** | 0 | 26 | 3 | **9** |  | 1 | 31 |
| 17/LHD | 26 | 51 | 0 | 35 | 5 | 6 |  | 1 |  |

RHD = right hemisphere damage; LHD = left hemisphere damage; MMSE = Mini Mental Scale Examination; Attention = Attentional matrices test; KF-NAP = Kessler Foundation Neglect Assessment Process; Raven = Raven test; MCST = Modified Card Sorting Test with scores for category (ability to find hidden rules) and perseveration errors (as a sign of impulsiveness). These tests are reported in raw scores. Memory: tests were different according to side of lesion, Rey’s Auditory Verbal Learning Test for RHD patients and Rey figure for LHD patients; these tests assess long-term memory, and data are reported with equivalent values (i.d., corrected tests’ scores by age and education, which range from 0 to 4 for both tests and permit comparison). Only LHD patients had an evaluation of language comprehension by Token test from Aachener Aphasie Test. Values under cut-off for each test are in bold (cut-off is calculated in accordance to each test’s practice, after correcting data for age, education and, if requested, gender).

***Results of rPCA on the computerized task data***

Table S2 reports the component loadings obtained from rPCA on the computerized task data. Note that rPC1 explained 41.9% of the variance, while rPC2 and rPC3 explained 23% of the variance each. High factor loadings, indicative of high contribution to the component, are highlighted in bold for visualization purposes. The first component (rPC1), on which we focus in the paper, loaded especially onto lateralized conditions (i.e., Contra- and Bi-lateral) but only under multitasking (i.e., both AVT and VDT).

**Table S2.** Component loadings from rPCA analysis

| **Loading** | **rPC1** | **rPC2** | **rPC3** |
| --- | --- | --- | --- |
| Bilateral Auditory Dual Task | **0.92** | 0.05 | 0.03 |
| Bilateral Single Task | 0.25 | -0.1 | **0.82** |
| Bilateral Visual Dual Task | **0.96** | 0.06 | -0.07 |
| Contralesional Auditory Dual Task | **0.81** | -0.02 | 0.22 |
| Contralesional Single Task | 0.39 | -0.16 | **0.64** |
| Contralesional Visual Dual Task | **0.84** | 0.11 | 0.11 |
| Ipsilesional Auditory Dual Task | -0.04 | **0.85** | 0.22 |
| Ipsilesional Single Task | -0.24 | **0.52** | **0.69** |
| Ipsilesional Visual Dual Task | 0.23 | **0.96** | -0.27 |

In bold higher coefficients, which mostly explain information about each rPC.

***Supplementary analysis: age-matched groups***

Cluster analysis was repeated after iteratively removing the youngest patients in C1 until the two groups were matched in terms of age. Figure S1 shows the results of the computerized task in the two age-matched groups.

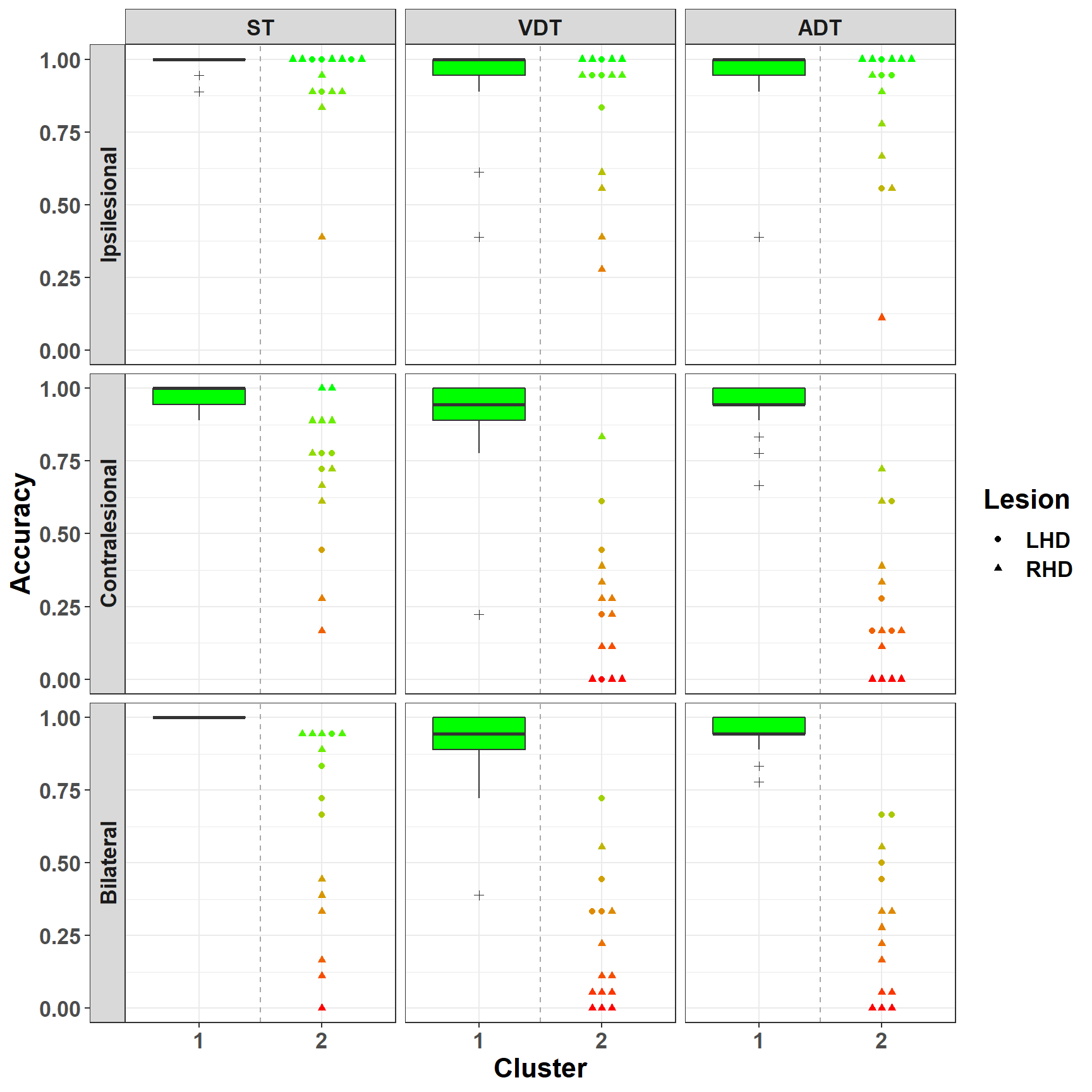

**Figure S1.** Data after removing the younger patients in C1 until reaching evidence that the two clusters (i.e., C1 e C2 from cluster analysis from the entire sample) were comparable in terms of age (BF10 <1). Accuracy in the computerized task  is depicted as a function of stimulus Type (i.e., Ipsi-, Contra-, or Bi-lateral presentation) and Load condition (i.e., single task, auditory-dual task, and visual-dual task).

**Neuroanatomical data**

***Overlays of lesions and structural disconnections***

Figure S2 shows overlays of lesions and structural disconnections normalized to the MNI space. The overlays show the most frequently damaged areas and tracts in patients (25 RHD patients, 15 LHD patients).

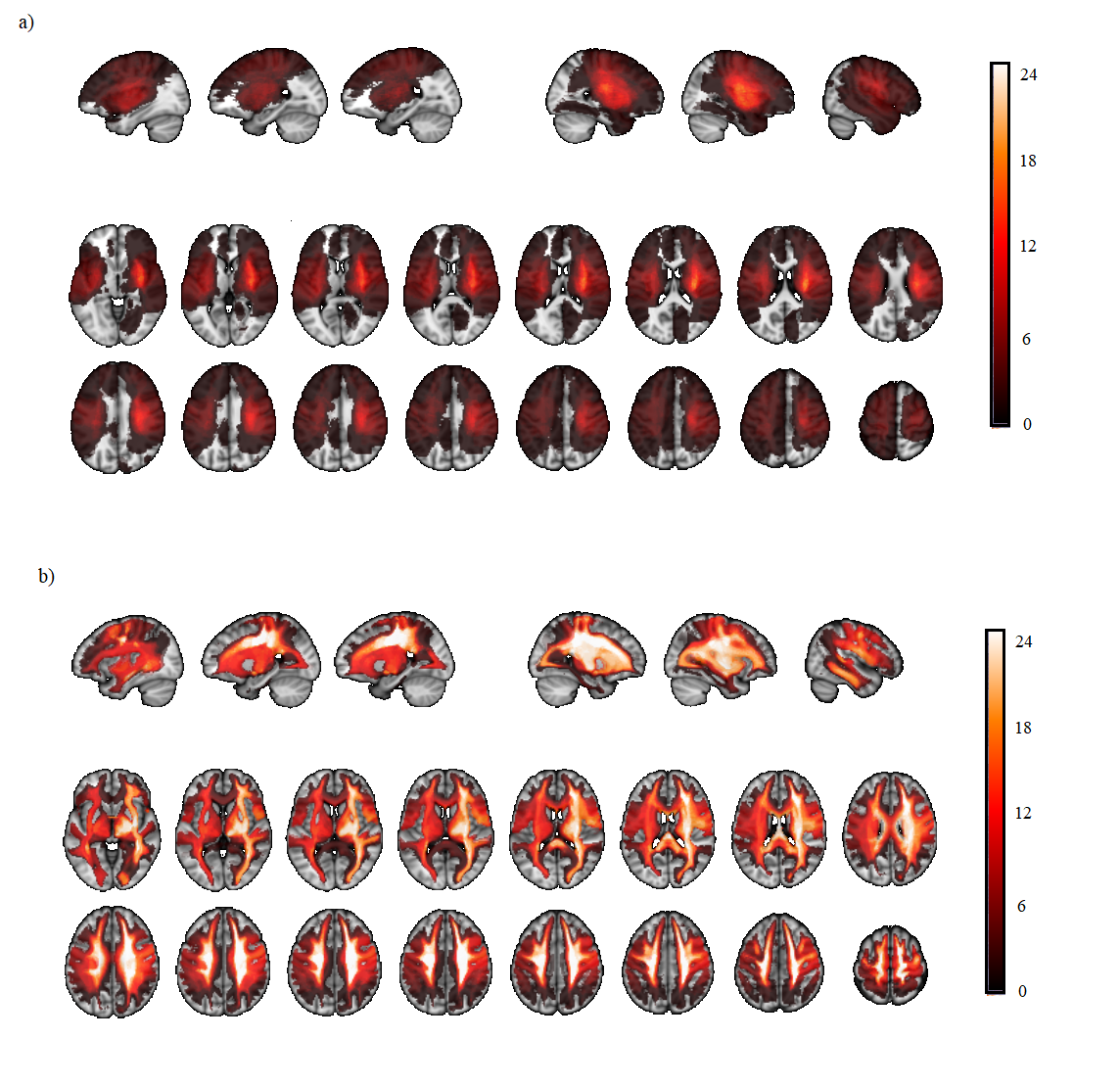

**Figure S2.** Overlay of a) lesions and b) disconnectomes on a standard MNI template. The color bar indicates the number of lesions overlapping across patients.
