## Supplementary material for "Susceptibility to multitasking in chronic stroke is associated to damage of the multiple demand system and leads to lateralized visuospatial deficits": Table 1

**Table 1.** Demographic, neurological and behavioral information for all patients.

| **Patient/**  **Group** | **Age** | **Gender** | **Education** | **Etiology** | **Onset**  **(months)** | **Lesion**  **(cm^3^)** | **BIT** |  |  | **Cluster** |
| --- | --- | --- | --- | --- | --- | --- | --- | --- | --- | --- |
| 1/RHD | 51 | M | 13 | I | 1.3 | 19.79 | 145 |  |  | 1 |
| 2/RHD | 49 | M | 8 | H | 3.3 | 19.59 | 143 |  |  | 1 |
| 3/RHD | 79 | M | 5 | I | 15.5 | 1.27 | 137 |  |  | 2 |
| 4/RHD | 56 | M | 8 | H | 3.5 | 68.19 | 141 |  |  | 1 |
| 5/RHD | 68 | M | 13 | I | 42.87 | - | 141 |  |  | 1 |
| 6/RHD | 75 | M | 13 | I | 3.53 | 10.72 | 139 |  |  | 2 |
| 7/RHD | 73 | M | 5 | I | 2.13 | 10.11 | 141 |  |  | 2 |
| 8/RHD | 65 | F | 5 | H | 99.2 | 9.51 | 146 |  |  | 1 |
| 9/RHD | 46 | F | 12 | H | 43.5 | 67.11 | 139 |  |  | 1 |
| 10/RHD | 47 | F | 13 | I | 23.27 | - | 140 |  |  | 1 |
| 11/RHD | 44 | M | 8 | H | 2.7 | 9.06 | 133 |  |  | 1 |
| 12/RHD | 76 | M | 5 | I | 4.3 | 33.73 | 137 |  |  | 1 |
| 13/RHD | 71 | M | 10 | H | 3.1 | 23.50 | 140 |  |  | 2 |
| 14/RHD | 64 | M | 16 | I | 5.3 | 169.67 | 133 |  |  | 2 |
| 15/RHD | 48 | M | 16 | I | 2.4 | 26.76 | 142 |  |  | 1 |
| 16/RHD | 51 | F | 8 | H | 2.5 | 6.79 | 144 |  |  | 1 |
| 17/RHD | 76 | M | 13 | I | 5.7 | 5.58 | 143 |  |  | 1 |
| 18/RHD | 75 | M | 13 | H | 12 | 18 | 142 |  |  | 1 |
| 19/RHD | 57 | F | 17 | H | 3.1 | 72.27 | 132 |  |  | 2 |
| 20/RHD | 82 | F | 13 | I | 3.3 | 26.39 | 133 |  |  | 2 |
| 21/RHD | 69 | M | 13 | H | 4.2 | 8.66 | 143 |  |  | 1 |
| 22/RHD | 75 | M | 5 | H | 1.9 | - | 138 |  |  | 2 |
| 23/RHD | 63 | M | 13 | I | 0.9 | 114.73 | 137 |  |  | 2 |
| 24/RHD | 68 | F | 8 | I | 1.5 | 7.70 | 138 |  |  | 1 |
| 25/RHD | 68 | M | 5 | I | 2.9 | - | 141 |  |  | 1 |
| 26/RHD | 49 | F | 8 | I | 3.1 | 222.40 | 141 |  |  | 1 |
| 27/RHD | 62 | M | 18 | I | 2 | 1.64 | 140 |  |  | 1 |
| 28/RHD | 72 | M | 13 | I | 74.8 | 329.96 | 136 |  |  | 2 |
| 29/RHD | 75 | M | 5 | I | 1.9 | 17.73 | 133 |  |  | 2 |
| 1/LHD | 42 | M | 18 | I | 24.6 | 37.05 | 142 |  |  | 1 |
| 2/LHD | 69 | M | 8 | I | 2.5 | - | 145 |  |  | 2 |
| 3/LHD | 45 | M | 13 | H | 10.4 | 49.19 | 144 |  |  | 1 |
| 4/LHD | 62 | M | 13 | I | 4.9 | 72.83 | 144 |  |  | 2 |
| 5/LHD | 61 | F | 5 | I | 1.2 | 10.11 | 145 |  |  | 1 |
| 6/LHD | 61 | M | 3 | H | 2.3 | 6.42 | 141 |  |  | 1 |
| 7/LHD | 61 | M | 8 | H | 3.6 | 4.21 | 144 |  |  | 1 |
| 8/LHD | 70 | M | 5 | H | 2.6 | 47.54 | 145 |  |  | 1 |
| 9/LHD | 47 | F | 8 | H | 4.4 | 17.81 | 142 |  |  | 1 |
| 10/LHD | 60 | M | 8 | H | 5.9 | 59.78 | 144 |  |  | 2 |
| 11/LHD | 64 | M | 18 | I | 2.3 | 137.13 | 143 |  |  | 1 |
| 12/LHD | 44 | M | 8 | H | 4 | 70.12 | 140 |  |  | 1 |
| 13/LHD | 77 | M | 8 | H | 2.8 | - | 140 |  |  | 1 |
| 14/LHD | 74 | F | 5 | H | 7.2 | 81.65 | 144 |  |  | 2 |
| 15/LHD | 37 | M | 13 | I | 1.5 | 126.67 | 145 |  |  | 1 |
| 16/LHD | 32 | F | 13 | I | 11.8 | 288.69 | 144 |  |  | 1 |
| 17/LHD | 69 | F | 11 | H | 2.7 | 16.12 | 140 |  |  | 1 |

RHD= Right Hemispheric damage, LHD= Left Hemispheric Damage, I= Ischemic, H= Hemorrhagic, Onset= Time from onset in months, Lesion= Size of lesion in cm**^3^**, BIT= Behavioral Inattention Test (conventional part), Cluster= results from clustering.
