## Supplementary material for "Susceptibility to multitasking in chronic stroke is associated to damage of the multiple demand system and leads to lateralized visuospatial deficits": Table 2

**Table 2.** Group differences and correlation between each variables and rPC1

|  |  | **Group differences** | | | | | **Correlation with rPC1** | | |
| --- | --- | --- | --- | --- | --- | --- | --- | --- | --- |
| **Measure** | **N(C1,C2)** | **C1**  **m±SD** | **C2**  **m±SD** | **t_(DF)_** | **p-value** | **r(c.i.)** | | **t_(DF)_** | **p-value** |
| Age | 46(31,15) | 57.35±12.63 | 70.07±7.31 | 4.32_(42.37)_ | **<0.001** | -0.46(-0.67,-0.2) | | 3.48_(44)_ | **0.001** |
| Lesion Volume | 40(27,13) | 49.47±69.56 | 79.79±88.38 | 1.09_(19.43)_ | 0.29 | -0.2(-0.48,0.12) | | 1.26_(38)_ | 0.22 |
| MMSE | 41(28,13) | 28.21±1.64 | 27.62±1.94 | 0.97_(20.32)_ | 0.35 | 0.14(-0.18,0.43) | | 0.88_(39)_ | 0.38 |
| BIT | 46(31,15) | 141.87±2.75 | 138.13±4.27 | 3.09_(19.82)_ | **0.005** | 0.54(0.3,0.72) | | 4.28_(44)_ | **<0.001** |
| Atten. matrices | 46(31, 15) | 44.23±10.09 | 37.13±9.8 | 2.28_(28.55)_ | **0.03** | 0.26(-0.03,0.51) | | 1.79_(44)_ | 0.08 |
| KF-NAP | 45(31,14) | 1.85±2.66 | 5.54±4.8 | 2.58_(15.63)_ | **0.02** | -0.58(-0.76,-0.33) | | 4.4_(38)_ | **<0.001** |
| Raven | 45(30, 14) | 30.81±3.62 | 27.57±5.37 | 2.05_(18.53)_ | 0.055 | 0.37(0.09,0.6) | | 2.64_(43)_ | **0.01** |
| MCST |  |  |  |  |  |  | |  |  |
| Category | 44(30, 14) | 4.77±1.3 | 3.57±2.17 | 1.9_(17.57)_ | 0.074 | 0.34(0.05,0.58) | | 2.33_(42)_ | **0.025** |
| Pers. errors | 40(27, 13) | 5.8±4.16 | 8.5±5.64 | 1.6_(19.88)_ | 0.126 | 0.26(-0.51,0.04) | | 1.77_(42)_ | 0.08 |

N= number of patients. m= mean and SD= Standard deviation.

MMSE= Mini-Mental State Examination, BIT= Behavioral Inattention Test, KF-NAP = Kessler Foundation Neglect Assessment Process, MCST= Modified Card Sorting Test

In bold significant results (p-values <0.05).
